# Neurodevelopmental timing and socio-cognitive development in a prosocial cooperatively breeding primate (*Callithrix jacchus*)

**DOI:** 10.1101/2023.12.01.569587

**Authors:** Paola Cerrito, Eduardo Gascon, Angela C. Roberts, Stephen J. Sawiak, Judith M. Burkart

## Abstract

Primate, and especially human, brain development is experience-dependent: it is shaped by the inputs received during critical periods. During early development, these inputs systematically differ between independently and cooperatively breeding species, because in cooperative breeders infants are interacting from birth with multiple caretakers and have to thrive in a richer and more challenging social environment. Here, we study the neurodevelopmental timing of the cooperatively breeding common marmoset and how it maps onto behavioral and developmental milestones. To obtain meaningful correlations of structure-function co-constructions, we combine behavioral, imaging (anatomical and functional) and neural tracing experiments. We focus on brain areas that are critically involved when observing conspecifics interacting with others and find that (i) these areas develop in clusters; (ii) these areas reach their maximum gray matter volume shortly after peak provisioning, when immatures are intensely provisioned by group members; (iii) the differentiation of these areas coincides with the period of intense negotiation between immatures and multiple adults over food, the birth of the next set of siblings, and the task of becoming a helper. Moreover, like in humans, differentiation is not fully completed at the age of first reproduction. In sum, we find that the developmental timing of social brain areas coincides with key social and developmental milestones in marmosets, and extends into early adulthood. This rich social input is likely critical for the emergence of the particularly strong prosociality and socio-cognitive skills of marmosets. Since humans are cooperative breeders too, these findings have strong implications for the evolution of human social cognition.

## Introduction

Strong social cognition and prosociality are, from a very young age, hallmarks of the human mind compared to the closest living relatives, the non-human great apes (Herrmann et al., 2007). Due to our peculiar life-history, characterized by early weaning and extensive allomaternal care very early in infancy (between 2 and 4 years of age) human development is embedded in a world filled with other individuals, including parents, siblings, and other family members. Thus, this is the context in which human toddlers’ strong social cognition and prosociality develops ((Hrdy & Burkart, 2020, 2022). It is this same period that is also the most important for the formation of the neural bases of higher-order social, emotional and communicative functions (Courchesne et al., 2007). Not surprisingly then, several independent lines of evidence, spanning neuroscience, pediatrics, primatology and psychiatry point to the fundamental role that the relative timing of brain development and social interactions have for the acquisition of social cognition and prosocial behaviors (Tottenham, 2020).

It is known that deviations in the normal developmental timing of the cortex can profoundly impact socio-cognitive skills and are one of the main factors linked to the occurrence of autism spectrum disorder (ASD) (Courchesne et al., 2007). Specifically, several studies have found that early brain overgrowth during the first years of life strongly correlates with ASD (e.g. Courchesne et al., 2005) and a meta-analysis of all published MRI data by 2005 revealed that the period of greatest brain enlargement in autism is during early childhood (Redcay & Courchesne, 2005), with about a 10% volume increase compared to controls during the first year of life. Hence, individuals affected by ASD present an accelerated early-life brain growth and achieve a final brain volume that is not different from that of controls, but they achieve it earlier than controls. Indeed, recent work with human brain organoids has confirmed the accelerated maturation of the cortex in the ASD phenotype (Paulsen et al., 2020; Urresti et al., 2021). Consequently, given this accelerated early-life brain development, fewer social inputs are available during the period when gray matter volume reduces to adult size and differentiates via experience-dependent pruning. Accelerated development of functional connectivity between certain brain areas (e.g. amygdala-prefrontal cortex) can also be a consequence of early life stress, which in turn can cause adverse physiological conditions such as increased anxiety and cortisol levels (Gee et al., 2013).

The importance of social inputs occurring during prolonged brain maturation and slow developmental pace has also been highlighted in the context of human evolutionary studies. The significant brain growth and development occurring postnatally in humans arguably allows the brain to be influenced by the social environment outside of the uterus to a greater extent than that seen in other great apes (Wilder & Semendeferi, 2022). Hawkes and Finlay (2018) show that in addition to weaning our infants earlier than expected (based on allometric scaling with other life-history variables), human neonates have an especially delayed neural development, which is likely correlated with the energetic trade-offs stemming from the large size and high caloric demand of our brain (Kuzawa et al., 2014). Additionally, we observe that in humans, compared to other great apes, myelination is much prolonged and continues well into adulthood (Miller et al., 2012).

Common marmosets (*Callithrix jacchus*) are cooperatively breeding platyrrhine monkeys. Like humans, they rely on extensive allomaternal care and share many life-history traits (e.g. short interbirth intervals and a hiatus between menarche and first reproduction) with humans (Hrdy, 2016). They also show remarkable prosociality (Burkart et al., 2007, 2014, much more than great apes, Burkart et al., 2014; Verspeek et al., 2022) and strong socio-cognitive abilities, which have been argued to correlate with cooperative breeding (Burkart & van Schaik, 2010; Hrdy & Burkart, 2022). However, the neurobiological features underlying the socio-cognitive abilities promoting prosocial behavior are poorly understood. Moreover, experimental research has shown that in common marmosets (hereafter: marmosets) there is a critical period for the development of social behaviors (Dettling et al., 2002); although the relationship between developmental timing of the brain and these early-life social interactions is poorly understood.

Given these similarities with humans, they are becoming an ever-more important model in neuroscience (Burkart & Finkenwirth, 2015; Miller et al., 2016; Oikonomidis et al., 2017; Samandra et al., 2022), and particularly in research investigating the neurobiological and neurodevelopmental bases of social cognition. As in humans, immature marmosets are surrounded and cared for by multiple caregivers from the first day on. The entire family is typically present during birth and oxytocin levels increase not only in mothers, but in all group members (Finkenwirth et al., 2016). Group members contribute significantly to carrying the infants and once infants start eating solid food, frequently share food with them. After a peak provisioning period, adults are increasingly less willing to share food with them (Guerreiro Martins et al., 2019; Rapaport, 2011; Sehner et al., 2022). During this period, intense and noisy negotiations over food are frequent, with immatures babbling and begging and adults eventually giving in – or not. Intriguingly, when doing so, immatures appear to take into account how willing individual adults are to share and will insist in more and longer attempts with adults who are generally less likely to refuse them. Soon after, immatures not only have to compete for attention and food with their twin sibling, but also with the next offspring that are born far before they themselves are independent because, like in humans, marmosets are weaned early and mothers have their next offspring soon after (Schultz-Darken et al., 2016). By now, the immatures still have not reached puberty; this only happens shortly before yet another set of younger siblings is born. Typically with these new arrivals the immatures start to act as helpers themselves, and thus face the developmental task of switching from being a recipient of help to becoming a provider of help and prosocial acts (Yamamoto, 1993). This is thus the developmental context in which marmoset socio-cognitive skills develop.

The goal of this study is to map the behavioral milestones specific to a cooperatively breeding primate to its region-specific brain development to better understand the social interactions in which infants engage during the differentiation period of brain regions differentially implicated in processing social stimuli. Our working hypothesis is that, like in humans, social interactions with several caregivers during this critical period profoundly contribute to the co-construction of the marmoset brain, the maturation of socially-related associative areas and, therefore, the emergence of prosocial behaviors. For that purpose, we sought to determine whether there is a relationship between the temporal profile of the developing marmoset brain and the early-life social interactions which may help explain their sophisticated socio-cognitive skills at adulthood.

To compare the timing of brain development to that of these behavioral milestones and developmental tasks, we focused on brain regions that in adult marmosets are selectively activated by the observation of social interactions between conspecifics, but not by multiple, but independently behaving marmosets, as identified by Cléry and colleagues (2021). We tested whether these brain regions share similar developmental trajectories based on the developmental patterns of regional gray matter volume. To potentially reveal a coordinated ontogenetic profile underlying the “tuning” of the social brain in marmosets, we then compared these neurodevelopmental patterns to longitudinal data of infant negotiations with caregivers in relation to food (as measured by the frequency of food begging).

We thus combined several types of data collected from marmosets in order to provide a unified picture of structural brain development alongside the development of social interactions between infants and multiple caregivers necessary to ensure survival (infant provisioning). These included structural magnetic resonance data (sMRI) of gray matter (GM) of 53 cortical areas and 16 subcortical nuclei acquired from a developmental cohort (aged 13 to 104 weeks) of 41 male and female marmosets (Sawiak et al., 2018); functional magnetic resonance data (fMRI) mapping the brain areas activated by the observation of social interactions in marmosets (Cléry et al., 2021); food sharing interactions in five family groups of marmosets including a total of 26 adults and 14 immatures (from 1 to 60 weeks of age, Guerreiro Martins et al. 2019), and cellular-resolution data of corticocortical connectivity in marmosets obtained via 143 retrograde tracer injections in 52 young adult marmosets of both sexes (Majka et al., 2020).

Overall, we make the following predictions:

P1: cortical regions that show significantly stronger activation during the observation of social interactions (Cléry et al. 2021) share similar structural neurodevelopmental profiles, which are distinct from those regions showing significantly stronger activation during the observation of non-social activities.

P2: brain regions (both cortical and subcortical) showing a significantly stronger activation during the observation of social interactions exhibit a protracted development, reaching their adult volume later than the other regions.

P3: the developmental trajectory of infant negotiations with caregivers in relation to food (as measured by the frequency of food begging) is more similar to that of brain regions responding more strongly to the observation of social interactions than to the other regions.

P4: functional connectivity is stronger between regions with similar developmental timing and response strength to the observation of either social or non-social behaviors, and weaker between regions with different developmental timing and response strength.

## Materials and Methods

### Materials

#### Cortical and subcortical structural developmental data

The data were collected in 41 male and female animals aged between 13 and 104 weeks, with each animal scanned at least twice using a Bruker PharmaScan 47/16 MRI system (Bruker, Inc., Ettlingen, Germany). Temporal milestones describing the structural developmental trajectories of the 69 brain regions considered have already been published (Sawiak et al., 2018). The data include 53 cortical areas and 16 subcortical nuclei. In the present work we accessed non-published, GM region-specific volumetric data, compiled across individuals using cubic spline models, following (Sawiak et al., 2018). Since both hemispheric asymmetry and sexual dimorphism were tested for and excluded (Sawiak et al., 2018) we use pooled male and female right and left hemisphere values.

#### Functional magnetic resonance (fMRI) data

These data were obtained from a published source (Cléry et al., 2021) and were acquired on male and female adult awake marmosets. Each animal was recorded while presented with video stimuli of social interactions (playing, grooming etc.) and non-social behaviors (eating, foraging, etc.), as well as to the scrambled versions of the same videos (four conditions in total). For each monkey, all four conditions were analyzed and compared, revealing the brain regions that show significantly stronger activation while observing the social condition comparing to all other three conditions (Cléry et al., 2021). These areas have been coded as “social” in our dataset. The areas showing instead a significantly stronger activation during the observation of non-social behaviors have been coded as “non-social” and all others as “neither” (Data S1).

#### Provisioning data

Infant provisioning patterns were recorded in five family groups, including a total of 14 immatures (aged 1 to 60 weeks) and 26 adults, representing both male and female breeders, and male and female helpers (older siblings) (Guerreiro Martins et al., 2019). Once a food item was given to an individual, it was recorded whether each food item was shared with the infant proactively, was shared with the infant in response to infant’s begging, or was refused to a begging infant. Hence, sharing could be either proactive, facilitated or resisted, while the response to begging could be either refusing or sharing. The total proportion of food proactively shared and begged was then compiled across infants and family groups for each point in time, providing a developmental trajectory of both. The trajectories of proactive, facilitated and resisted sharing are very similar among each other (Guerreiro Martins et al., 2019). Conversely, the proportion of refused food increased as the one of shared food decreased. The trajectory of food begging reflects the complex negotiation dynamic that goes from helplessly receiving, to increasingly more interacting with caregivers in order to elicit a desired outcome (food). These interactions and negotiations with multiple caregivers are a behavioral peculiarity of cooperatively breeding animals (such as humans and marmosets) which entail infant provisioning by a variety of individuals as a consequence, also, of prolonged post-weaning dependence (Hrdy, 2016).

#### Corticocortical connectivity data

These data have been published in an open-access form (Majka et al., 2020) and are available on the following web platform: https://www.marmosetbrain.org. We used the full extrinsic fraction of labeled neurons (FLNe) connectivity matrix which was obtained via 143 retrograde tracer injections in 52 young adult marmosets of both sexes. The matrix reports the results of injections centered in 55 target areas of the marmoset cortex, including subdivisions of prefrontal, premotor, superior temporal, parietal, and occipital complexes. The data represents the weighed and directed connectivity matrix based on the results of injections of monosynaptic retrograde fluorescent tracer injections. The rows and columns of the matrix represent individual cortical areas. The targets are the injected areas while the sources indicate the areas in which the projections originate. The intrinsic connections are excluded from our analysis.

## Methods

### P1 – do ‘social’ areas share similar trajectories?

To obtain a finite number of meaningfully different developmental trajectories of the cortical areas, we followed Sawiak and colleagues (2018) and identified six clusters: the visual and cingulate cluster; the somatomotor cluster; the auditory-visual cluster; the orbitofrontal, dorsolateral and ventromedial prefrontal cluster; the ventrolateral prefrontal cortex (PFC), polar, operculum and insula cluster; the lateral and inferior temporal lobe cluster. Increasing cluster numbers resulted in the further subdivision of the prefrontal cortex, which is the area presenting the largest heterogeneity of developmental trajectories. We then matched the sMRI and fMRI data, which resulted in 32 regions for which we had both types of data (Data S1). This clusterization, combined with the matching of the sMRI and fMRI data, allowed us to test whether regions with similar response to social or non-social visual stimuli present similar developmental trajectories (i.e. belong to same developmental clusters), and therefore whether developmental timing and socio-cognitive function are correlated. Subcortical nuclei are excluded from this clusterization.

### P2 – do “social” areas have protracted development?

To quantify and compare the developmental timing of the different regions we defined four different milestones: A) the age at maximum volume; B) the age at the first descending inflexion point (the first local maxima of the first derivative); C) the age at maximum rate of volume decline; D) the age at the last ascending inflexion point (the last local minima of the first derivative). Three of these milestones (A, B and C) are standard ones that have been previously used (e.g. Sawiak et al., 2018), while we decided to introduce a novel one (D) to capture the decrease in rate of decline visible towards the end of the trajectories. For a visual representation and example of the four milestones, see Figures S1 and S2. Using these four milestones we defined three temporal ranges: age at maximum volume to age at first descending inflexion point (A to B); age at maximum volume to age at fastest decline (A to C); age at maximum volume to age at the last ascending inflexion point (A to D). Finally, we proceeded to compute Bonferroni-corrected pairwise t-test for each of the four milestones and three ranges, to assess whether they were significantly different between “social” and “non-social” regions.

### P3 - are regional trajectories of “social” brain areas similar to social feeding behaviors?

Based on the results of the analyses carried out to verify P2, we selected the milestones and ranges that distinguish the developmental trajectories of social regions from non-social ones and computed them also for the developmental trajectories of proactive food sharing and food begging. We then compared the values (in weeks) obtained for the provisioning data to those of the neurodevelopmental trajectories to test our prediction that interindividual negotiations (such as those occurring when infants beg caregivers for food) follow a similar developmental pattern to that of social brain regions.

### P4 – do regions with similar GM developmental profiles and fMRI response to stimuli of social interactions have stronger functional connectivity?

First, we matched the corticocortical connectivity data with the GM neurodevelopmental data, which resulted in a dataset of 1400 connections encompassing 28 target and 51 source regions (Data S2). We also matched the connectivity data with the fMRI data and included only the connections for which fMRI data is present for both the target and the source. This resulted in 494 connections between 19 target and 27 source regions (Data S3). To test whether connection strength differs between regions with similar response to social or non-social visual stimuli we performed Wilcoxon-rank sum tests and adjusted the results for multiple-hypothesis testing. We used non-parametric tests because of the non-normal distribution of the data. We also performed the same tests on the data grouped according to the six developmental clusters. To test whether there is a correlation between developmental timing and connection strength across regions, for each connection we computed the average age at peak rate of volume decline between the source and target region, and the absolute difference in age at peak rate of volume decline between the source and target region. We then built four models with fraction of extrinsic labeled neurons (FLNe) as response variable and either average age at peak rate of volume decline (avg_age_max_rate) or absolute difference in age at peak rate of volume decline (d_age_max_rate) as explanatory variables, and fMRI response type in source and target regions as either: i) additive variable: lm1 = *FLNe ∼ avg_age_max_rate + Source_to_Target* and lm2 = *FLNe ∼ d_age_max_rate + Source_to_Target* or ii) interaction term: lm3 = *FLNe ∼ avg_age_max_rate * Source_to_Target* and lm2 = *FLNe ∼ d_age_max_rate * Source_to_Target. We compared each pair of models (with Source_to_Target as either additive or interaction term) using AIC (Akaike, 1973)*.

## Results

Overall, our results support our predictions and show that: (i) brain areas that respond significantly more strongly to visual stimuli of social interactions belong to the same developmental clusters to the exclusion of regions that respond more strongly to visual stimuli of individuals acting alone; that (ii) social clusters have an extended developmental trajectory, reaching adult-like volumes later and after a period of plateau; and that (iii) the ontogenetic trajectory of food negotiations matches quite well that of “social” brain regions. However, we did not find support for stronger corticocortical connections between areas with stronger responses to visual stimuli of social interactions: social regions did not appear to form a strongly connected network.

### P1 – do ‘social’ areas share similar trajectories?

There is a biunivocal correspondence between those brain regions that were differentially activated (as measured via fMRI) by either social interactions or solitary behaviors (Figure 1A) and neurodevelopmental clusters (Figure 1B). Specifically, social interactions activated regions that fell into three developmental clusters; cortical areas 1 and 2, 3a, 4ab, 6DC and 8C cluster together; areas 14C, 25, 46D and 9 belong to another cluster; areas 10, 45, 47L and the proisocortical motor region belong to yet another cluster. Conversely, the seven cortical areas that responded more strongly to the observation of solitary behaviors, fell into the other three of the six developmental clusters: visual areas 1, 2 and 6 form one cluster; visual areas 3 and 4 form another cluster; and temporal area TE1/2 and the ventral temporal lobe form another cluster. As per our prediction, a given developmental cluster contains areas responding more strongly only to one of the two conditions.

**Figure 1.**
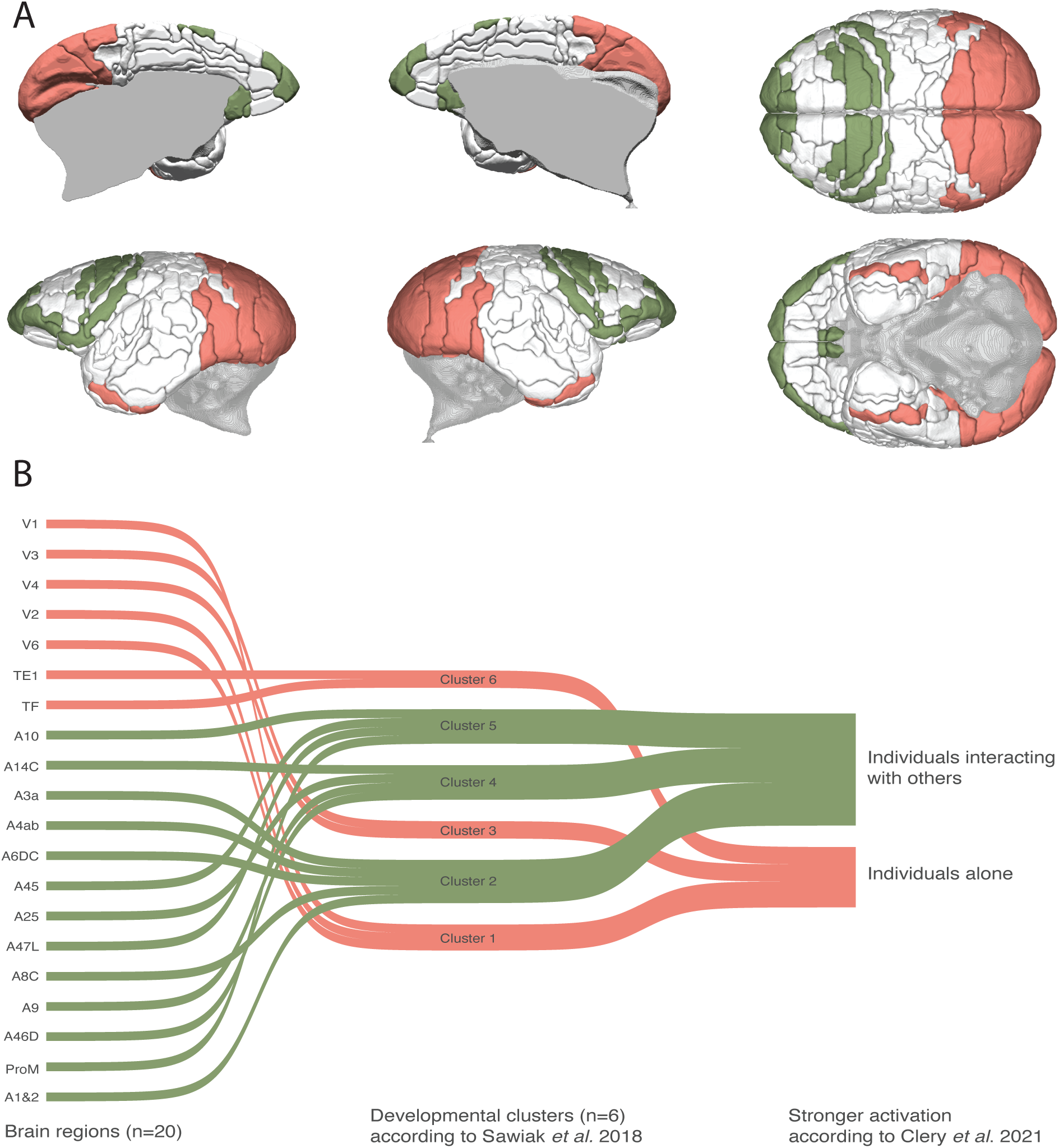
**A**: Location of the 20 cortical brain regions that are significantly more active (fMRI data) when observing individuals interacting with conspecifics (green), or individuals acting alone (pink). From top left moving clockwise: medial views of the left and right hemispheres, superior view, inferior view, lateral views of the right and left hemispheres. **B**: Sankey diagram of these same brain regions and their relationship to the six cortical clusters reported in Sawiak et al (2018): the visual and cingulate cluster (1); the somatomotor cluster (2); the auditory-visual cluster (3); the orbitofrontal, dorsolateral and ventromedial prefrontal cluster (4); the ventrolateral prefrontal cortex (PFC), polar, operculum and insula cluster (5); the lateral and inferior temporal lobe cluster (6).

### P2 – do “social” areas have protracted developmental?

The results of the Bonferroni-corrected pairwise t-tests for each of the four milestones and three ranges (Table 1, Figure 2) indicate that the age at the fastest rate of volume decline is the only milestone that differs significantly between regions that respond more strongly to visual stimuli of social interaction vs. solitary behaviors. Moreover, it is the temporal range between the age at maximum volume and the age at fastest rate of volume decline that distinguishes regions with differing activation to the two types of stimuli. The duration of the volumetric “plateau”, measured as time from the age at maximum volume to the age at which there is the first descending inflexion point (Figure S1) differs significantly (p=0.011) between brain areas that respond more strongly to visual stimuli of social interactions than to those of solitary behavior.

**Figure 2.**
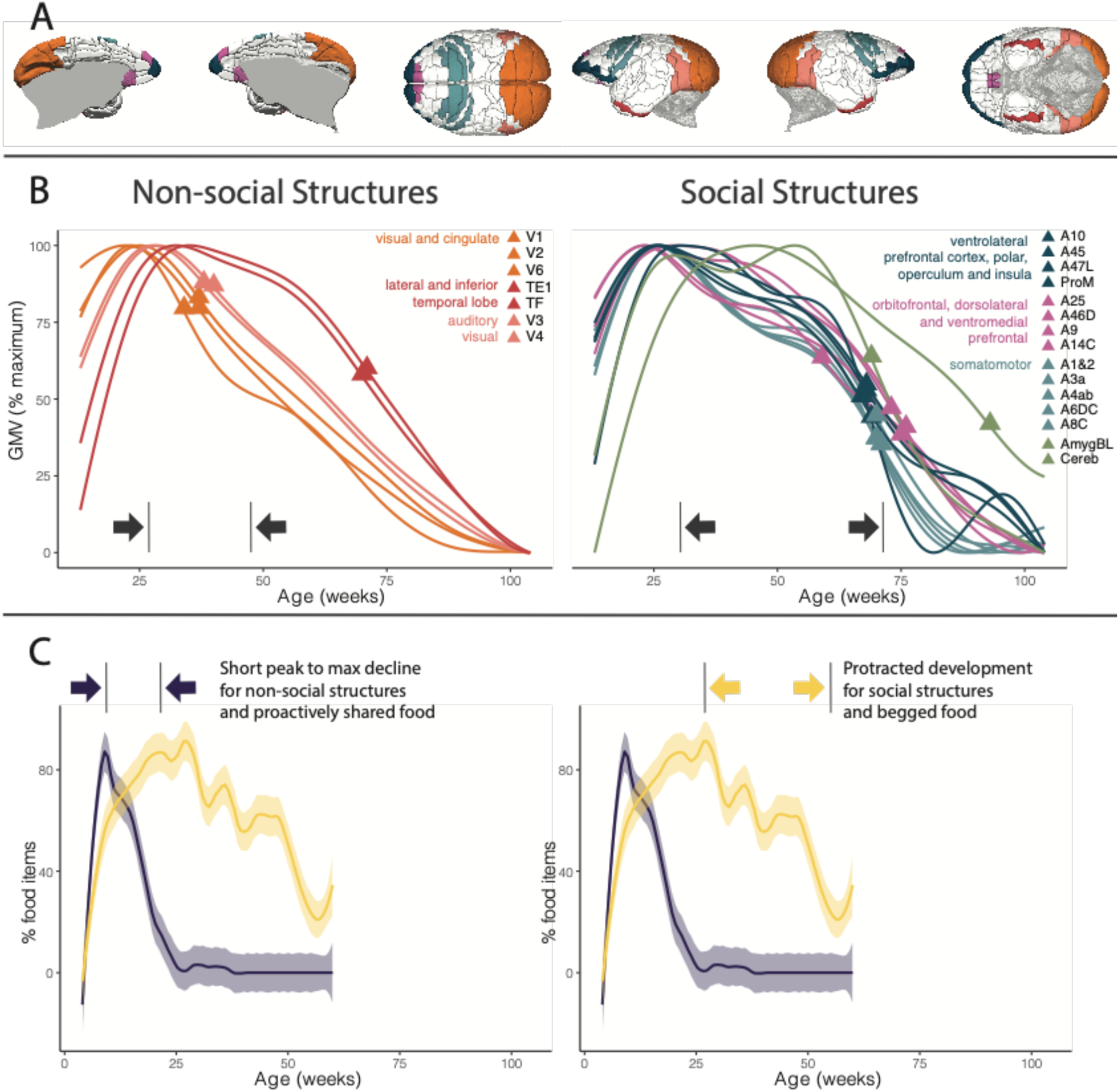
**A**: Location of the 20 cortical regions analyzed. Colors represent developmental clusters and match the colors in panel B. **B**: Developmental profiles of the “non-social” (left) and “social” (right) regions. The triangle is placed on point C (age fastest rate of decline). The colors represent the six developmental clusters of cortical regions, while the sub-cortical regions (basolateral nucleus of the amygdala and cerebellum) are represented in green. The x-axis represents time (expressed in weeks from birth), while the y-axis represents relative volume (absolute values are not reported graphically as there is tremendous variance between regions and would therefore be difficult to visualize all together). **C**: In blue, the proportion of food that is proactively shared, in yellow the proportion of trials for which infant begging occurs (regardless of the outcome of begging), the shaded areas indicate the 95% C.I. The ranges reported in the four plots indicate the time between developmental milestones A (maximum value) and C (maximum rate of decline).

**Table 1:**
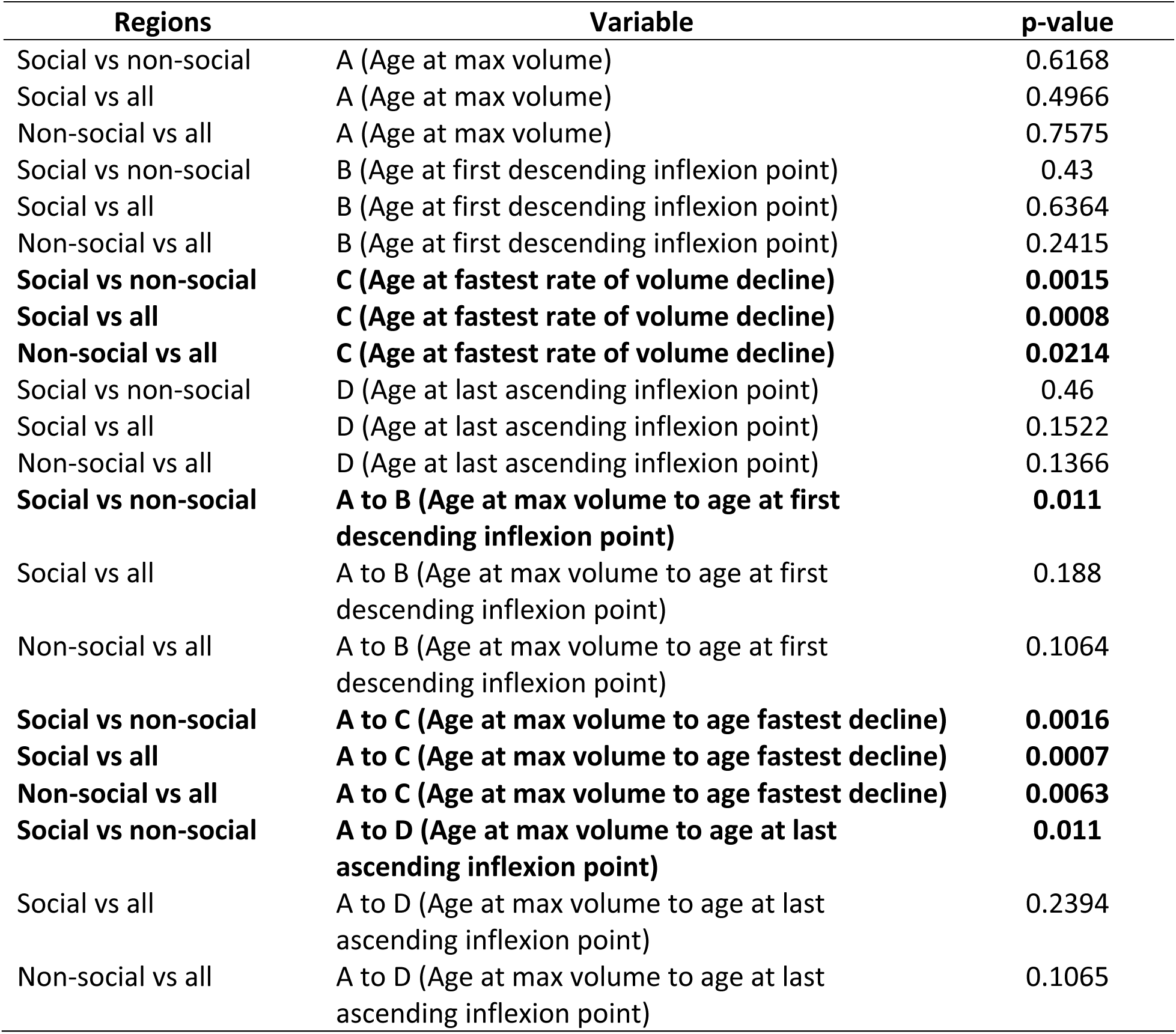
Results of Bonferroni-corrected pairwise t-tests comparing milestone and ranges between brain regions that have significantly stronger activation in response to either social or non-social stimuli. Bolded results are significant at p < 0.05.

P3 - are regional trajectories of “social” brain areas similar to those of social feeding behaviors? The results of the comparisons between developmental trajectories of brain regions and food proactively shared or begged support our prediction. The age at fastest decline is much later for the proportion of food begged (58 weeks) than for the proportion of proactively shared food (25 weeks). Moreover, the range between the peak and the age at fastest decline is also larger for the proportion of food begged (31 weeks) than for the proportion of food shared (17 weeks). The ontogenetic trajectory of food begging parallels that of the brain regions with stronger activation in response to visual stimuli of social interactions, while that of proactively shared food parallels that of the brain regions with stronger activation in response to stimuli of solitary behaviors (Figure S3, Table 2). The ontogenetic trajectories of the two patterns of food provisioning (proactively sharing and begging) parallel those of brain regions (respectively, non-social and social) but anticipate them (Figure 2).

**Table 2:** For the ontogenetic trajectories of brain regions average values and standard deviations are reported, while for the ontogenetic trajectories of food sharing the single value is reported (all values are in weeks).

| Variable | Average<br>(*Value) | SD |
| --- | --- | --- |
| C Social regions | 71.3 | 7 |
| C Non-Social regions | 46.7 | 16.3 |
| C Proportion begged | *58 |  |
| C proportion proactively shared | *25 |  |
| A to C Social regions | 42.2 | 8.25 |
| A to C Non-Social regions | 19 | 12.3 |
| A to C Proportion begged | *31 |  |
| A to C proportion proactively shared | *17 |  |

### P4 – do regions with similar GM developmental profiles and fMRI response to stimuli of social interactions have stronger functional connectivity?

Our results (Table 3) indicate that connectivity strength is significantly different (and greater) between regions that both respond more strongly to non-social stimuli, rather than between those responding more strongly to social stimuli. The regions that respond more strongly to non-social stimuli are also the ones with faster developmental trajectories, which cluster in groups 1, 3 and 6. The results when the same analysis is performed on those regions grouped by developmental cluster, rather than in response to different stimuli, are reported in the supplementary materials (Table S1, Figure S4). Models with *Source_to_Target* as additive term had lower AIC scores than the corresponding model with *Source_to_Target* as interaction term. The results of the two models with additive term (Tables S2 and S3) indicate that there is a weak correlation between similarity in developmental timing and connectivity strength (for both models: R^2^ = 0.2; p<0.0001). We used the Akaike Information Criterion (AIC) (Akaike, 1973) to compare the two models, and the one having as response variable the absolute difference in age at maximum rate of volume decline is the best one. Finally, in both models, the intercept is significantly different (p<0.0001) only between regions that both respond more strongly to non-social stimuli, thus confirming the results of the Wilcoxon-rank sum tests (Figure S5).

**Table 3:** results of the pairwise Wilcoxon-rank sum tests (p-values are corrected for multiple hypothesis testing) on the connectivity strength between brain regions with different activation strength to different types of stimuli. Values reported as 0 are < 0.00000001.

|  | Neither<br>to<br>Neither | Neither<br>to<br>Nonsocial | Neither<br>to<br>Social | Nonsocial<br>to<br>Neither | Nonsocial<br>to<br>Nonsocial | Nonsocial<br>to<br>Social | Social to<br>Neither | Social to<br>Nonsocial |
| --- | --- | --- | --- | --- | --- | --- | --- | --- |
| Neither to<br>Nonsocial | 1 | NA | NA | NA | NA | NA | NA | NA |
| Neither to<br>Social | 0.44866 | 1 | NA | NA | NA | NA | NA | NA |
| Nonsocial to<br>Neither | 0.70614 | 1 | 1 | NA | NA | NA | NA | NA |
| Nonsocial to<br>Nonsocial | **0.01302** | **0.00104** | **0.00007** | **0.00072** | NA | NA | NA | NA |
| Nonsocial to<br>Social | **0.00001** | **0.00286** | **0.00045** | **0.00503** | **0** | NA | NA | NA |
| Social to<br>Neither | 1 | 1 | 1 | 1 | **0.00049** | **0.00001** | NA | NA |
| Social to<br>Nonsocial | **0.00155** | 0.44866 | 0.37016 | 0.70293 | **0** | 0.44866 | **0.00695** | NA |
| Social to<br>Social | 1 | 1 | 1 | 1 | **0.00093** | **0.00049** | 1 | 0.21406 |

## Discussion

Our findings reveal that those brain regions more strongly activated when viewing visual social interactions have different ontogenetic trajectories from those that are not (P1), indicating that developmental timing and function are correlated in marmosets. Specifically, while the various brain regions do not differ significantly with respect to the age at which they reach their maximum GM volume, those involved in processing social interactions maintain the maximum volume significantly longer, before decreasing in size and reaching their adult value (P2). Since GM volumetric decline is a consequence of synaptic pruning and intra-cortical myelination (Paus, 2005), it functionally corresponds to a decrease in developmental plasticity.

Of all the regions analyzed, the one with the latest peak in volumetric decline and therefore prolonged plasticity is the basolateral nucleus of the amygdala. This nucleus is implicated in encoding emotional events with reference to their particular sensory-specific features (Balleine & Killcross, 2006) and has undergone convergent evolution in its volumetric patterns in cooperatively breeding species including marmosets and humans (Cerrito & Burkart, 2023). The slowest developing cortical area is 46D. It is located in the dorsolateral prefrontal cortex, together with areas A10 and A9, which are also slow in developing and respond more strongly to social stimuli. In humans, the dorsolateral prefrontal cortex has been shown to be implicated in theory of mind (Xi et al., 2011) and in the suppression of selfish behaviors (van den Bos et al., 2011). The next slowest developing brain region is A14C, located in the ventromedial prefrontal cortex together with areas 45, 47L and the proisocortical motor region (ProM). A14C is also known to be heavily implicated in social cognition (Delgado et al., 2016), including joint attention (Williams et al., 2005), facial emotion recognition, theory-of-mind ability and processing self-relevant information in humans (Hiser & Koenigs, 2018). Interestingly, a longitudinal study in humans affected by ASD has shown that differential activation during a temporal discontinuing task in the ventromedial prefrontal cortex, including A14C, and cerebellum in these individuals was associated with abnormal functional brain maturation (Murphy et al., 2017). Another study has shown a decrease in effective connectivity from the temporal pole (another late-maturing brain region) to the ventromedial prefrontal cortex, and overall lower activation in the latter (Rolls et al., 2020).

Our findings on regional developmental timing in marmosets are similar to those recently described in humans (Bethlehem et al., 2022). Both species have similar temporal ranges, in relation to developmental milestone, in the ages at which the different regions reach their maximum volume (2 to 10 years in humans; roughly corresponding to the 22 to 77 weeks in marmosets) (Charvet et al., 2023). The same study on humans also highlighted an important similarity to our findings in marmosets: primary sensory regions showed earlier post-peak declines, whereas fronto-temporal association cortical areas showed later post-peak declines.

It is during the period of peak provisioning by non-maternal family members that gray matter volume is dramatically increasing (Figure 3), potentially highlighting the fundamental role played by provisioning in providing the nutritional and energetic needs of the developing brain.

**Figure 3:**
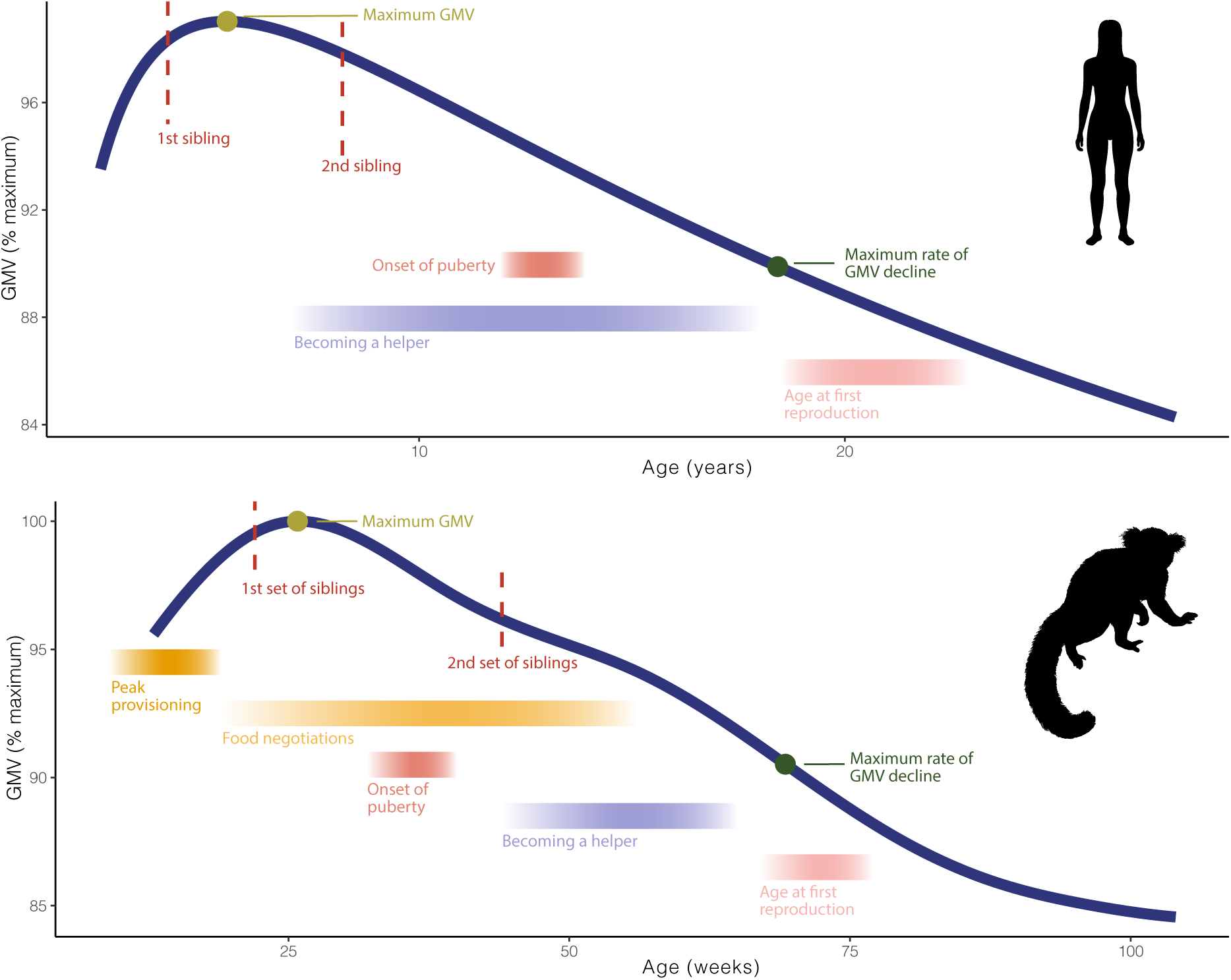
Comparison between cortical gray matter volume (GMV) trajectories and behavioral and developmental milestones in humans (top) and common marmosets (bottom). Comparative GMV data for humans is from Bethlehem et al., 2022. Data for behavioral and developmental milestones for humans refer to hunter-gatherer populations and are compiled from several sources (Henry et al., 2005; Kramer et al., 2017; Meehan et al., 2013). Behavioral and developmental data for marmosets are obtained from the following sources: Guerreiro Martins et al., 2019; Schultz-Darken et al., 2016.

The ontogenetic trajectory of regions more strongly activated by the observation of social interactions resembles that of food begging (which is inherently socially interactive), but not of proactively shared food, which does not require active communicative participation from the side of the infant (P3). The proportion of proactively shared food peaks at 8 weeks of age (ranging from week 7-20), a time during which breeders and alloparental caregivers proactively offer to infants almost 40% of the food items. Conversely, infant begging peaks at 27 weeks of age, with values remaining high (above 50%) until 50 weeks. Infant begging is a form of interaction and communication that requires an active participation of the infant (unlike proactive sharing), who must learn to engage and elicit care most efficiently from a plurality of individuals (parents and helpers). Anticipating the current motivation and intention of these individuals and adjusting their own solicitation behavior accordingly are important means by which immatures can reach high efficiency in achieving their goal.

Research on humans has shown that the elicitation of care from non-maternal caregivers entails the mobilization of more regulatory efforts than during the interactions with mothers (Feldman & Klein, 2003). Furthermore, comparative work on primates (Cerrito & DeCasien, 2021) indicates that in species practicing alloparental care there is greater control of the facial musculature and facial expressions, which enable non-vocal communication between infants and caregivers. These findings align well with recent work (Dureux et al., 2023; Schaeffer et al., 2020) that used fMRI to identify face-selective patches in the marmoset brain and found differential activation in subcortical regions and in the anterior cingulate and lateral prefrontal cortices while animals viewed socially relevant videos of marmoset faces. This differential activation was observed in many of the brain regions that are also the slowest to reach the age at maximum volume decline, such as the amygdala (102 weeks of age), the cerebellum (92.6 weeks), A32 (82.5 weeks) and A8 (70.1 weeks) which are located in the anterior cingulate, and A47L (68.3 weeks) which is located in the lateral prefrontal cortex. This combined evidence suggests that facial expressions are not only important for infant-alloparent communication, but that the brain regions employed to process these expressions have a prolonged plasticity compared to other brain regions (average age at fastest decline across all regions is 65.1 weeks), and that this prolonged ontogenetic trajectory maps onto that of infant-caregiver interactions in relation to food begging.

Limitations of this study are mainly in regard to sample sizes. First, while the fMRI data used in our work (Cléry et al., 2021) represents the most cutting-edge development and updated data of this type for marmosets, it is nevertheless collected on a small sample of subjects: three adult marmosets (one female, two males). However, other recent work (Schaeffer et al., 2020; Zanini et al., 2023) on marmosets employing fMRI to differentiate brain regions that are activated by visual stimuli depicting an intact action vs. its phase-scrambled version corroborate and replicate the results obtained by Cléry and colleagues (2021). Further work with an increased number of individuals representing several age groups would provide a valuable validation of our results and the possibility to assess how the structural development and the functional response of brain regions responding more actively to stimuli of social interactions map onto each other. Furthermore, due to slight differences in the parcellation of the cortical areas between the structural MRI and the functional MRI data (e.g. area 19 in sMRI data and area 19M in fMRI data), it was only possible to match 32 regions. Greater homology in the parcellation used would allow for an increased sample size in regions for which structural and functional data is both available. Ultimately, studies such as this one would benefit tremendously from collecting all the different types of data (neuroanatomical, functional and behavioral) from the same individuals. This is largely limited by the different research questions that are motivating each lab to collect a specific type of data, and by the facilities available to each lab. Ideally, future research could consider conducting longitudinal studies in which behavioral, anatomical and physiological measures are acquired for animals living in as “natural-like” conditions as possible.

Despite the above limitations, several lines of evidence highlight the distinct developmental trajectories of brain regions that display a significantly stronger response to visually presented social interactions compared to those of individuals acting alone. Specifically, regions implicated in the evaluation of social interactions have prolonged neurodevelopmental time periods. Furthermore, we show that this same neurodevelopmental trajectory mirrors, with a temporal shift of about 20 weeks, that of infant food begging, a form of care elicitation and infant-adult interaction that is necessary to ensure infant survival in cooperatively breeding species. More broadly, our results are in line with recent work (Ash et al., 2023) showing that in marmosets the critical period for the acquisition of cognitive control is between 39 and 65 weeks of age, falling within the temporal period from age at maximum GM volume and the age at fastest GM volume decline (Figure 3). This temporal range is also the one during which individuals transition from being a recipient of help to becoming a helper (Yamamoto, 1993), which arguably requires inhibitory control (i.e. to share rather than consume food items) (Feistner & Chamove, 1986). Our results are further supported by previous findings (Quah et al., 2022) showing that, in marmosets, family size has an effect on the age at which brain regions mature, with individuals belonging to larger families (and therefore having to integrate a larger number of different social interactions) having protracted neurodevelopment compared to those living in smaller families.

Finally, our current work underscores the striking similarity in the relationship between brain developmental patterns and behavioral milestones in marmosets and humans (Figure 3). In both species, perhaps as a consequence of the short interbirth intervals, immatures become helpers and become physiologically capable of reproduction while GMV volumes are still drastically changing, and even before GMV reaches its maximum rate of decline. Additionally, both species experience a hiatus between the onset of puberty and the age at which first reproduction actually occurs. This is likely because prior social experience with being a helper is necessary to carry out a successful reproductive event. Indeed, reproduction in callitrichids often fails if individuals have not been helpers first (Smith, 2005).

Phenotypic convergences between distantly related species as a consequences of changes in developmental timing are all but surprising. Indeed, across mammals, both allomaternal behavior and developmental timing are phylogenetically labile compared to most other traits (Blomberg et al., 2003; Cerrito & Spear, 2022). Hence, heterochrony (i.e. a difference in the timing, rate, or duration of a developmental process) can likely cause the phenotypic convergence observed in marmosets and humans, two species whose evolutionary trajectories diverged ∼42MYA (Upham et al., 2019) but that in many life-history aspects resemble each other more than each one resembles its sister taxon. Moreover, it is known that small differences in timing can result in large phenotypic changes, such that, for example, heterochronic divergence can cause paedamorphosis (i.e. the retention of an ancestral juvenile trait into adulthood) due to neoteny (Fenlon, 2022). Hence, changes in the timing of life history events, in relation to the relatively phylogenetically conserved neurodevelopmental schedule (Workman et al., 2013) can cause convergence in the adult phenotype. Future comparative studies of the neurodevelopmental and behavioral trajectories of independently-breeding, closely related taxa are the next fundamental step to effectively understand how the interplay between anatomical and behavioral timing shapes prosocial behavior.

While life-history and cognitive convergences between humans and marmosets have been abundantly documented, here we provide the first comparative evidence of what is potentially the developmental trajectory underlying the emergence of the adult phenotype (we address the mechanistic and ontogenetic aspect of the question, *sensu* Tinbergen, 1963). We argue that it is a delayed and prolonged brain development occurring during a time in which infants actively engage in social interactions and negotiations that allows for the emergence of prosociality: the brain needs time with alloparents and selective pressures to interact with them, in order to be tuned towards prosociality.

## Contributions

PC conceived the study and carried out the data analysis. PC and JMB designed the study, interpreted the results, and wrote the first draft of the manuscript. ACR and SJS collected the structural MRI data. JMB, ACR, SJS and EGG supervised the work and contributed to the final version of the manuscript.

## Ethics Approvals

Neuroimaging procedures were carried out in accordance with the UK Animals (Scientific Procedures) Act 1986 as amended in 2012, under project licences 80/2225 and 70/7618. In addition, the University of Cambridge Animal Welfare and Ethical Review Body (AWERB) provided ethical approval of the project licence.

