## Supplemental Materials for "Neurodevelopmental timing and socio-cognitive development in a prosocial cooperatively breeding primate (*Callithrix jacchus*)"

### Supplementary Tables

| term | estimate | std.error | statistic | p.value |
|---|---|---|---|---|
| **(Intercept)** | **0.019** | **0.009** | **2.102** | **0.036** |
| **Connectivity_S$d_age_max_rate** | **-0.001** | **0.000** | **-3.338** | **0.001** |
| Connectivity_S$Source_to_TargetNeither_to_Nonsocial | -0.002 | 0.012 | -0.184 | 0.854 |
| Connectivity_S$Source_to_TargetNeither_to_Social | -0.001 | 0.010 | -0.140 | 0.889 |
| Connectivity_S$Source_to_TargetNonsocial_to_Neither | -0.002 | 0.012 | -0.214 | 0.831 |
| **Connectivity_S$Source_to_TargetNonsocial_to_Nonsocial** | **0.096** | **0.013** | **7.461** | **0.000** |
| Connectivity_S$Source_to_TargetNonsocial_to_Social | -0.004 | 0.010 | -0.433 | 0.665 |
| Connectivity_S$Source_to_TargetSocial_to_Neither | -0.005 | 0.010 | -0.461 | 0.645 |
| Connectivity_S$Source_to_TargetSocial_to_Nonsocial | -0.001 | 0.011 | -0.096 | 0.924 |
| Connectivity_S$Source_to_TargetSocial_to_Social | 0.006 | 0.010 | 0.574 | 0.566 |

*Table S1: Results of the FLNe ~ d_age_max_rate + Source_to_Target model.*

| term | estimate | std.error | statistic | p.value |
|---|---|---|---|---|
| **(Intercept)** | **0.027** | **0.018** | **1.500** | **0.134** |
| Connectivity_S$avg_age_max_rate | 0.000 | 0.000 | -1.240 | 0.215 |
| Connectivity_S$Source_to_TargetNeither_to_Nonsocial | -0.005 | 0.013 | -0.405 | 0.686 |
| Connectivity_S$Source_to_TargetNeither_to_Social | 0.004 | 0.010 | 0.350 | 0.727 |
| Connectivity_S$Source_to_TargetNonsocial_to_Neither | -0.003 | 0.012 | -0.255 | 0.799 |
| **Connectivity_S$Source_to_TargetNonsocial_to_Nonsocial** | **0.096** | **0.013** | **7.123** | **0.000** |
| Connectivity_S$Source_to_TargetNonsocial_to_Social | -0.005 | 0.010 | -0.493 | 0.622 |
| Connectivity_S$Source_to_TargetSocial_to_Neither | 0.001 | 0.011 | 0.087 | 0.930 |
| Connectivity_S$Source_to_TargetSocial_to_Nonsocial | -0.008 | 0.011 | -0.750 | 0.454 |
| Connectivity_S$Source_to_TargetSocial_to_Social | 0.020 | 0.010 | 1.972 | 0.049 |

*Table S2: Results of the FLNe ~ avg_age_max_rate + Source_to_Target model.*

### Supplementary Figures

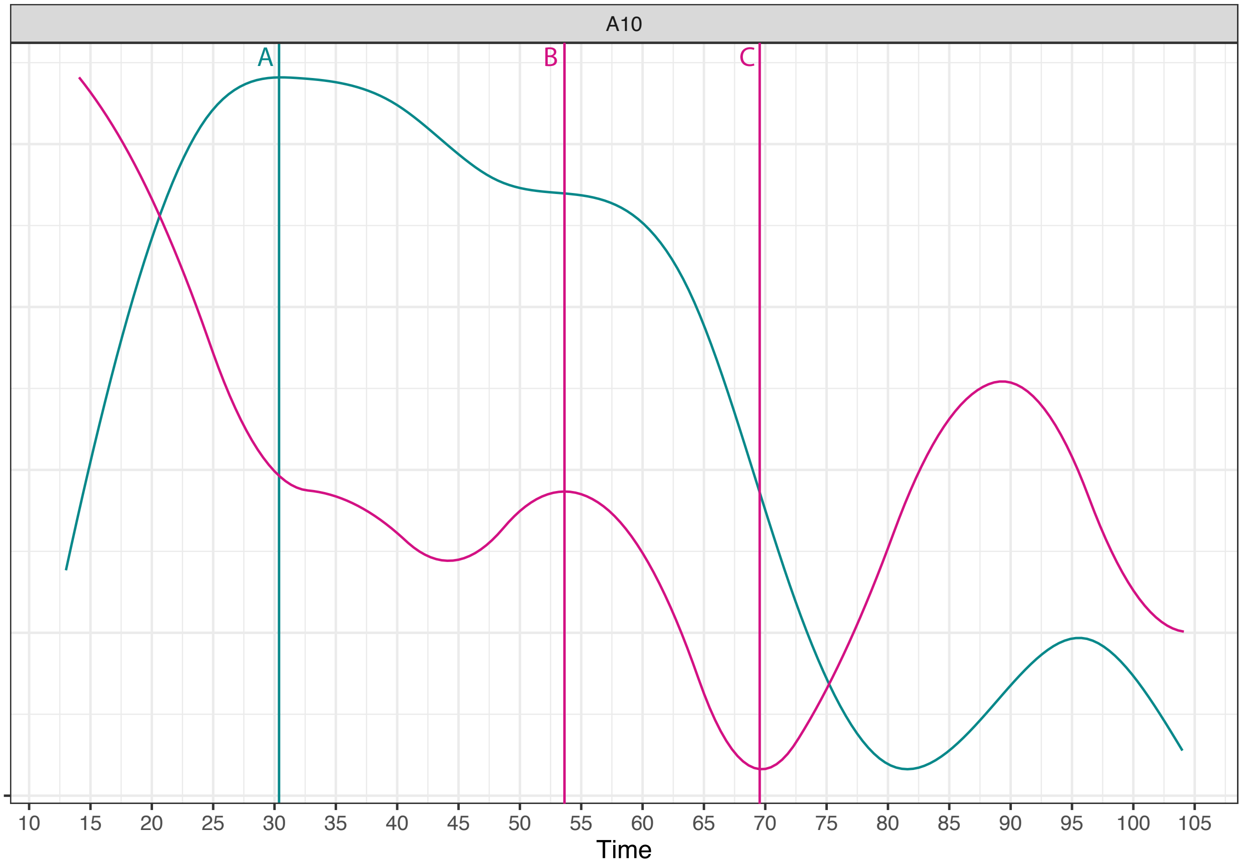


*Figure S1: Green: volume changes over time. Pink: First derivative of green curve. Y-axis is different for the pink and green curves, but not relevant here. A = age at maximum volume; B = Age at first descending inflexion point (the first local maxima of the first derivative); C = age at maximum rate of decline (global minima of the first derivative).*

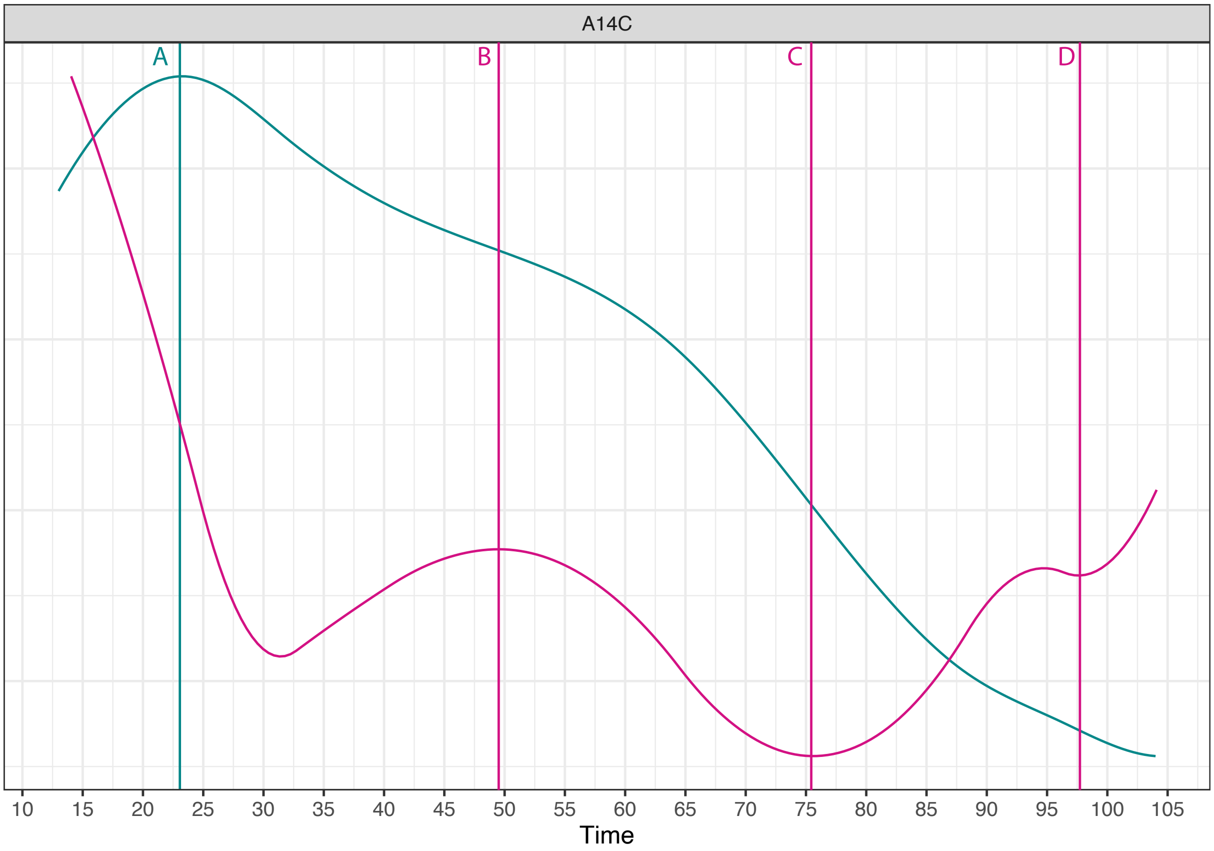


*Figure S2: Green: volume changes over time. Pink: First derivative of green curve. Y-axis is different for the pink and green curves, but not relevant here. A = age at maximum volume; B = Age at first descending inflexion point (the first local maxima of the first derivative); C = age at maximum rate of decline (global minima of the first derivative); D = age at last local minima of the first derivative.*

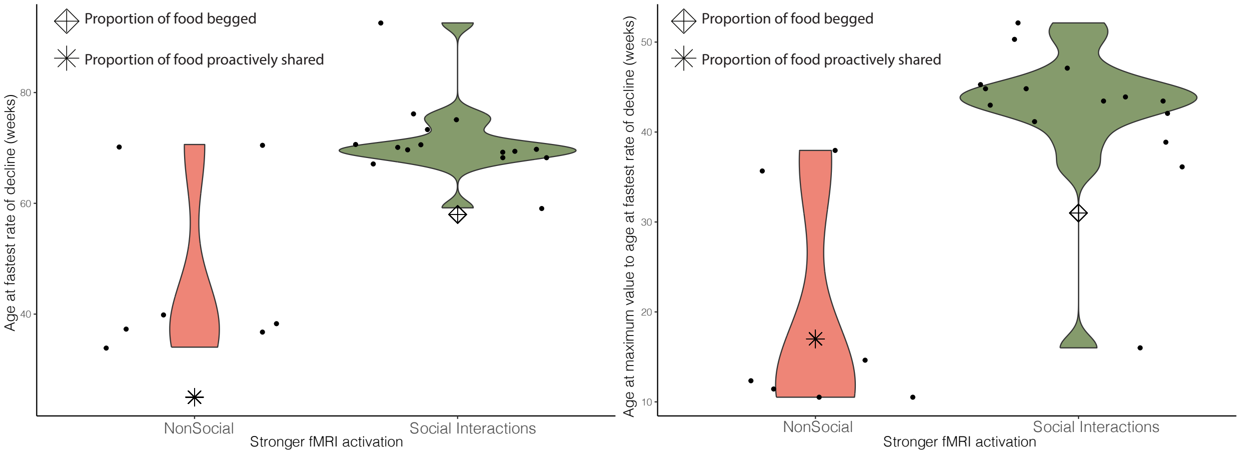


*Figure S3: For social regions (i.e. those activated when observing social interactions, green), the fastest decline of gray matter volume occurs later than for non-social regions (left), and the plateau from the maximum value to the fastest decline is longer (right). These same milestones and ranges are also shown for the ontogenetic trajectories of proactive food sharing (star) and food begging and negotiation (diamond). Individual dots represent individual brain regions.*

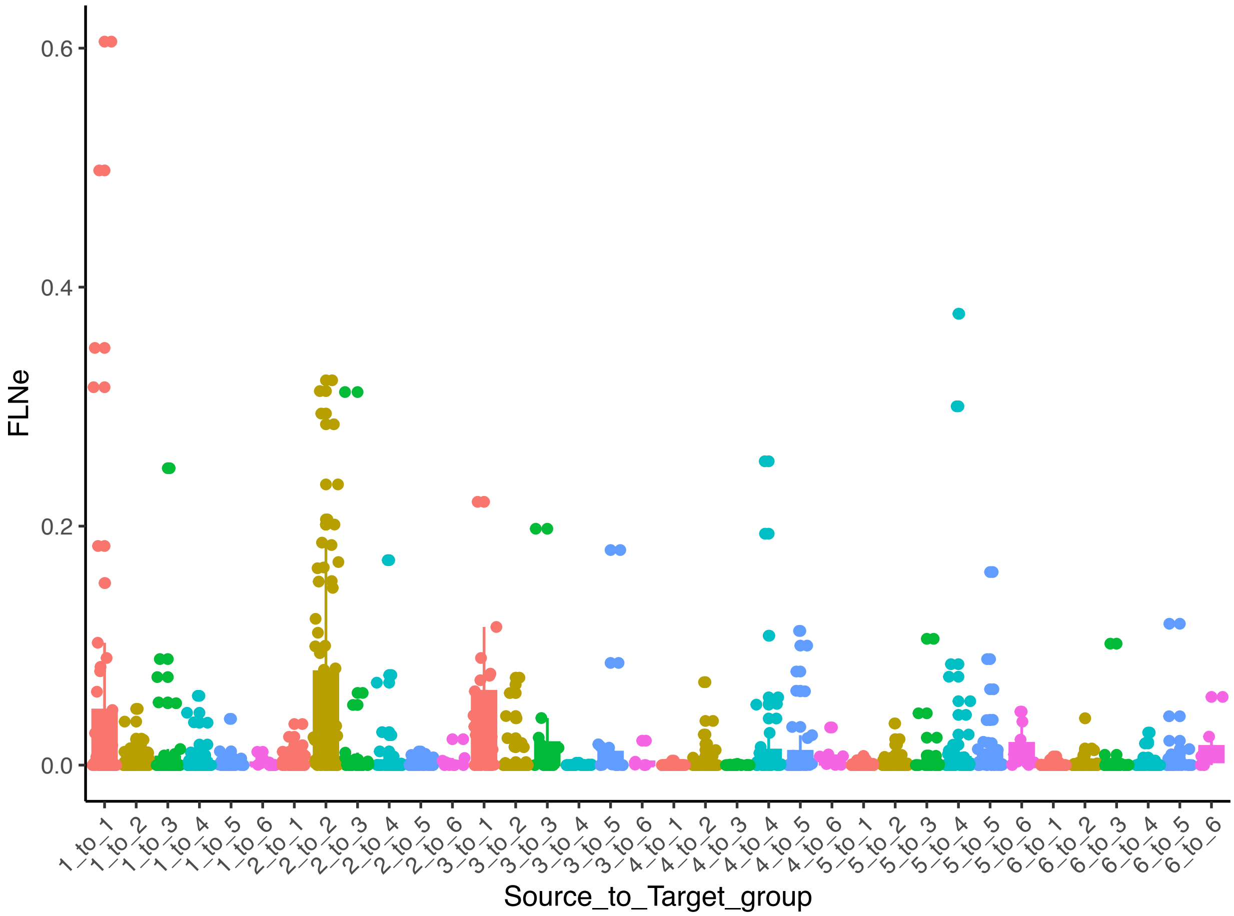


*Figure S4: fraction of extrinsic labeled neurons (FLN2) colored according to the developmental cluster of the target regions. The numbers of both the source and target regions refer to the developmental clusters, as reported in (Sawiak et al., 2018) and used throughout the manuscript. Developmental clusters for each of the regions are reported in Data S1.*

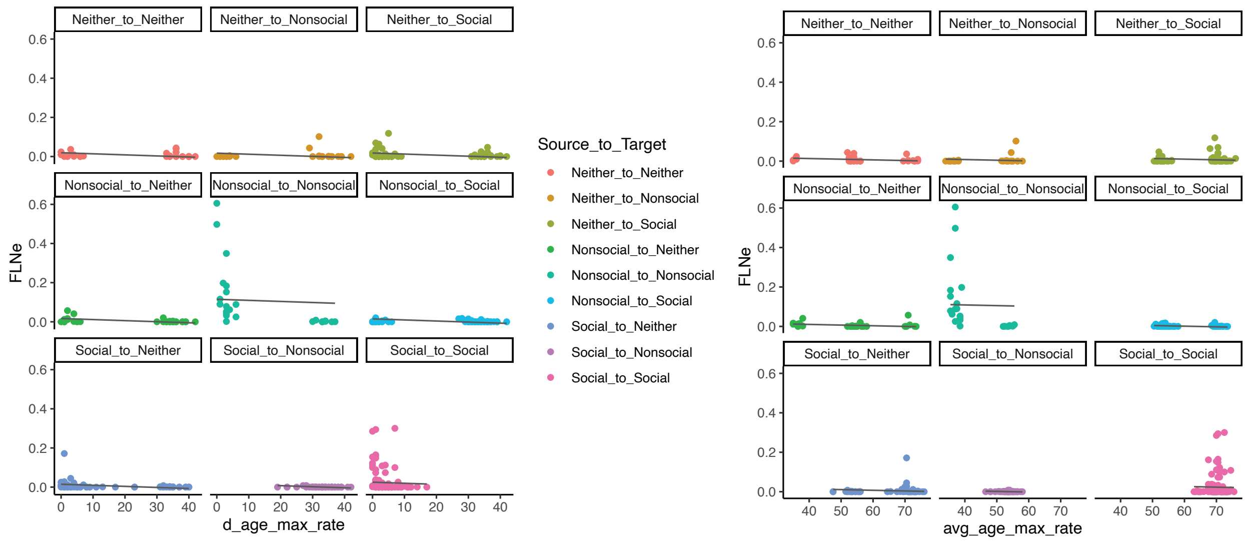


*Figure S5: Plots showing the correlation between fraction of extrinsic labeled neurons (FLNe) and two different measures of similarity in developmental timing (expressed in weeks). Left: absolute difference in the age at which the maximum rate of GM volumetric decline is reached. Right: average age at which the maximum rate of GM volumetric decline is reached. The regression lines are those for FLNe ~ d_age_max_rate + Source_to_Target (left) and FLNe ~ avg_age_max_rate + Source_to_Target (right). In both models, the intercepts are significantly different only for non-social to non-social regions. The overall R-squared is 0.2 for both models.*
